## Supplementary Figures for "Context-dependent selectivity to natural scenes in the retina"

**Title**

Affiliations:
1 Institut de la Vision, Sorbonne Université, INSERM, CNRS, Paris
2 Laboratoire de physique de l'Ecole normale supérieure, CNRS, PSL University, Sorbonne University, and University of Paris, Paris, France
3 Institute of Computer Science and Campus Institute Data Science, University of Göttingen, Germany

*: equal contributions
†: corresponding authors

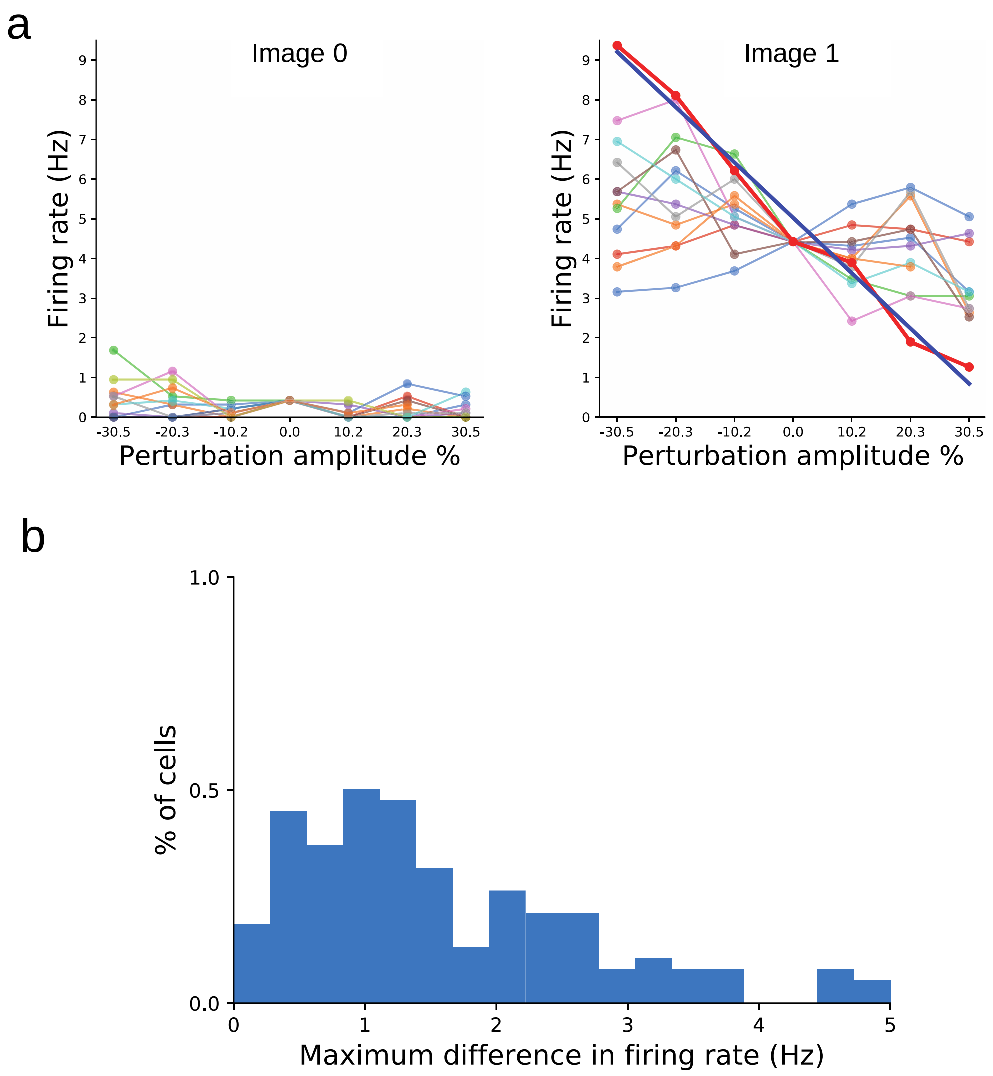


**Supplementary Figure 1. Calibration for the perturbative approach amplitude.**

(A) Four natural images were selected as reference, and 12 perturbation patterns were added to them using 6 different amplitudes (an amplitude of 100% would correspond to adding an amplitude equal to the mean luminance of the image). Each perturbed image was displayed to a mouse retina 25 times. All presentations were interleaved. The two panels show the 12 response curves for two images and each perturbation for an example ganglion cell, as a function of the perturbation amplitude. The response was estimated by counting spikes in a time window from 30 ms to 350 ms after image presentation. Here, the ganglion cell responds mostly to Image 1. For each curve, we fitted a linear regression and calculated the slope. We picked the perturbation having the highest slope, across all images, for each cell. We used this maximal slope to estimate, for a given perturbation amplitude, the difference between the firing rate evoked by the “maximal” perturbation, and the response to the image alone.

(B) Distribution of the maximal change in firing rate as estimated above, across many ganglion cells (n=140), for a perturbation amplitude of 25%. This amplitude gave us an average change of 1.7 Hz, corresponding to an average change in the spike count of 0.5 in the time window used to estimate the response. The selection criteria for the value of 32 was that we obtained at least 0.5 mean spike variation with respect to the non-perturbed image for each perturbed image presentation. A value of 12.5% was selected for salamander in the same manner.

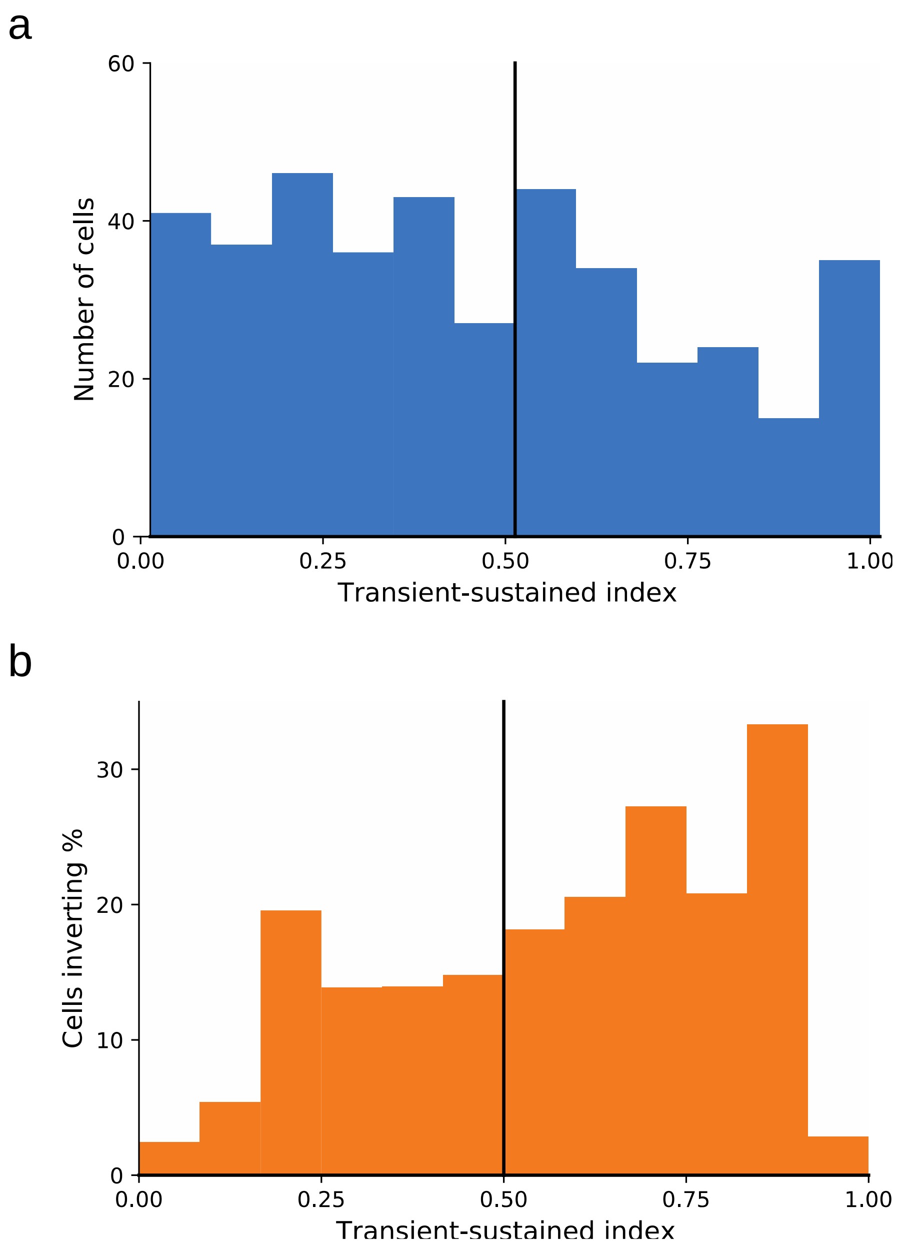


**Supplementary Figure 2. Polarity inverting ganglion cells are more transient.**
(A) Distribution of the Transient-Sustained index computed for all mouse retinal ganglion cells. We estimated the index from ganglion responses to a full-field flash. The early response E was estimated as the number of spikes emitted from 0 to 0.8 s after the beginning of the stimulus. The late part of the response L was estimated as the average spike count from 0.8 to 1.6 s after the onset of the stimulus. The index is defined as: (E-L)/(E+L). If the index is 1 it means that all the spikes occurred in the early window, thus indicating a transient response. On the contrary, if the index is near 0, it means that there are as many spikes in the late window as in the early one, indicating a sustained response.
(B) For each of the bins in A, we calculated the percentage of cells that showed polarity inversion during our perturbative approach.

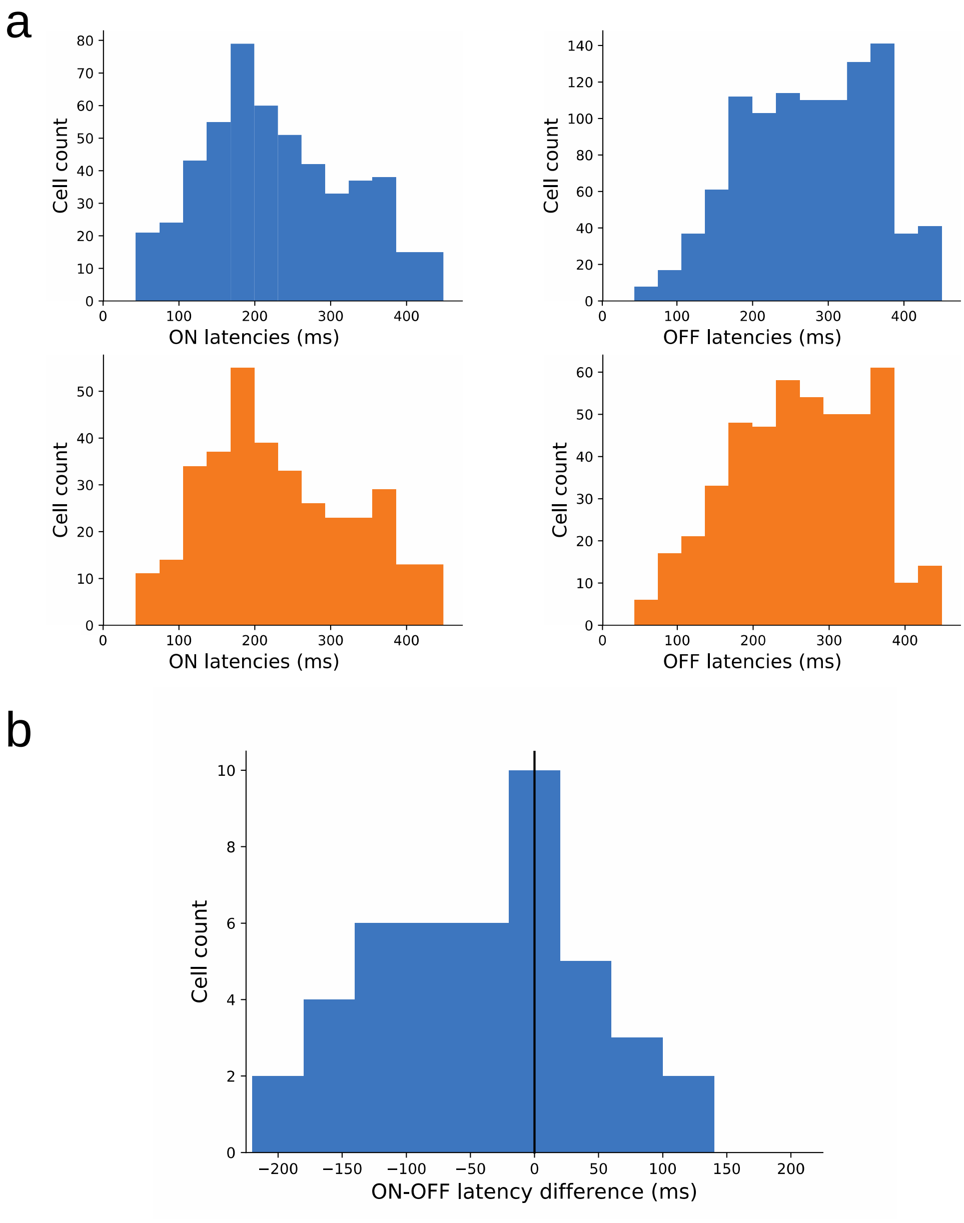


**Supplementary Figure 3. Latency difference between ON an OFF LSTAs for the polarity inverting cells.**

(A) We calculated the ON latency (left column) and OFF latency (right column) for all mouse retinal ganglion cells (top row, blue) for the perturbed images where an LSTA was detected. To smooth the average responses, each spike after a stimulus was replaced by a Gaussian centered at the time of the spike with a standard deviation of 20 ms, and then all the curves were averaged to obtain the average response to the perturbations. Then, we looked for the peak of the resulting curve in a window starting from 30 ms to 450 ms after the perturbed image presentation. We did the same for the polarity inverting retinal ganglion cells (bottom row, orange), with no noticeable difference with the whole population.

(B) Difference in latency between the responses associated to ON vs OFF LSTAs for the retinal ganglion cells showing polarity inversion. The peak of the distribution is at 0, but the long tails of the distribution of latencies go from -220ms to +140 ms.
